# A Pooled CRISPR Screen Protocol for Comparative Pathway Analysis Across Multiple Stressors

**DOI:** 10.64898/2026.09.07.749902

**Authors:** Truc T. Losier, Karyn E. King, James Taylor, Maxime W.C. Rousseaux, Ryan C. Russell

**Author notes:** Lead contact: 451 Smyth Rd, Ottawa, Ontario, K1H 8M5, Canada (Ryan C. Russell).

## Abstract

High content CRISPR screens have become a leading method for the identification of new cellular pathways, proteins functions, and drug targets. However, as the complexity of these screens increase, so does the possibility of artifacts. Here, we describe a refined CRISPR screening protocol that can compare acute cellular responses across multiple cellular treatments and timepoints, while reducing false positives by up to 40% compared to conventional screening approaches. This is achieved through the use of a fluorescence-based reporter line responsive to multiple stressors, as well as a fixation step to preserve acute response differences and avoid sorting-related false positives. We recently performed a version of this workflow to compare kinases regulating different types of selective autophagy, including starvation-induced autophagy, ERphagy, and pexophagy. Beyond autophagy research, this screen approach can be implemented for any fluorescence-based screen where preservation of acute cellular responses is important. Here we present a complete workflow of this optimized screening protocol using our selective autophagy screen as a specific example throughout. Implementation of these strategies should allow researchers to increase the number of stressors tested within a single screen without sacrificing cell health, reliability, or statistical power.

## Introduction

In recent years, high-throughput genetic screening technologies, including RNAi- and CRISPR/Cas9-based approaches, have transformed functional genomics, enabling systematic identification of gene regulators in mammalian cells. Regulators of autophagy in particular, have been successfully identified using RNAi and CRISPR screens^1–12^. Autophagy is a highly conserved catabolic process that enables cells to degrade and recycle cytoplasmic components to maintain homeostasis and adapt to stress^13,14^. While bulk autophagy non-selectively engulfs cytosolic material, selective autophagy removes specific cargo such as damaged organelles, protein aggregates, and pathogens through defined receptor-mediated processes^15–20^. Uncovering specific regulators of selective autophagy is critical for the understanding of disease pathologies and the development of increasingly targeted therapeutic approaches^21,22^. Despite a need to determine distinct regulators of selective autophagy, most autophagy screens have been limited to studying a single pathway or stress condition. This is largely because a comparative analysis is technically challenging due to differences in reporter sensitivity across conditions, asynchronous processing of cells across varying treatment times, and time constraints during cell sorting.

To overcome these challenges, we developed a CRISPR screening protocol that allows for the parallel analysis of multiple stress-induced pathways in the same batch of reporter cells. Modifications to basic workflows include the use of a single reporter acting as a readout for multiple stressors, and a pre-sort fixation step to preserve cellular state. We previously used this workflow to perform a kinome-wide screen, which successfully identified regulators of five different autophagy pathways^23^. For this screen, we used a DsRed-IRES-GFP-p62 reporter capable of interacting with multiple types of ubiquitinated cargo and fixed cells with 4% paraformaldehyde (PFA) prior to sorting to capture acute signaling events and diminish sorting artifacts.

Using our published autophagy screen as a practical example, we outline the optimization steps used to effectively design and execute a multi-condition screen. Although demonstrated here in the context of autophagy and selective autophagy, the methodological framework is broadly applicable to multi-condition functional genomics, drug target discovery, and pathway mapping. This integrated approach addresses several key challenges in comparative pooled screens, offering a reproducible, adaptable strategy for dissecting complex cellular processes under diverse experimental perturbations.

## Materials

### Cell culture

A critical consideration when designing pooled genetic screens is the selection of an appropriate cell line. The ideal cell line will exhibit robust and stable reporter expression, is sensitive to the biological processes under investigation, maintains good cell health and viability, and is amenable to efficient transduction or transfection using either viral or non-viral delivery methods. In this study, we chose HEK293A cells (Thermo Fisher Scientific#R70507) because they activate autophagy in response to a wide variety of stressors^23–26^ HEK293A cells were cultured in DMEM supplemented with 10% bovine calf serum (VWR Life Science Seradigm). All cells were routinely screened for mycoplasma.

### Amino acid free media

The recipe for amino acid starvation media is described in Table 1.

**Table 1.** Recipe for amino acid free media.

| Chemical | Working concentration (mM) |
| --- | --- |
| Calcium Chloride (CaCl<sub>2</sub>) | 1.8 |
| Ferric Nitrate Solution (Fe(NO<sub>3</sub>)<sub>3</sub>·9H<sub>2</sub>O, stored at 4°C) | 0.000248 |
| Magnesium Sulfate (MgSO<sub>4</sub>·7H<sub>2</sub>O) | 0.814 |
| Potassium Chloride (KCl) | 5.33 |
| Sodium Bicarbonate (NaHCO<sub>3</sub>) | 44.05 |
| Sodium Chloride (NaCl) | 81.9 |
| Sodium Phosphate monobasic (NaH<sub>2</sub>PO<sub>4</sub>·H<sub>2</sub>O) | 0.906 |
| D-Glucose (Dextrose) | 25 |
| HEPES | 25.03 |
| Phenol Red (optional) | 0.0399 |

1. Listed chemicals are added into 800 ml of distilled H_2_O and the solution is stirred until completely dissolved.
2. pH is adjusted to 7.4.
3. 20 mL of 100X MEM vitamins (VWR#45000-702) is added so that the final concentration is 2X.
4. The total volume is brought up to 1 L.
5. The solution is filtered through 0.22 µm Stericup (Thermo Fisher Scientific #SCGPU11RE), aliquoted into 50 ml tubes, and stored at 4°C.

### Plasmids

pLenti-DsRed-IRES-eGFP vector is from Addgene (Cat#92194). Brunello human kinome CRISPR knockout library was obtained from Addgene (Cat#1000000083)^27^. To generate lentiviruses, lentiviral packaging plasmids psPAX2 (Addgene#12260) and pMD2.G (Addgene#12259) are transfected in HEK293T cells^28,29^ along with plasmid carrying either DsRed-IRES-GFP-p62 or human kinome CRISPR knockout library.

### Flow cytometry and Sorting

The intensity of the fluorescent reporter can be analyzed using flow cytometry. We detected DsRed and GFP-p62 signal using BD FACSCelesta flow cytometer equipped with a blue (488 nm) laser and a yellow green (561 nm) laser. Following flow cytometry analysis using FlowJo software, cells were sorted into low and high GFP:DsRed populations using a Sony Biotechnology SH800 sorter.

### Other equipment

Tissue culture hoods and incubators are required for the maintenance and treatment of reporter cell lines. Other necessary equipment includes a vortex, cell counter, PCR machine, and high-speed centrifuge, and NGS sequencer. Access to core facilities or services with next generation sequencing capacity is also essential.

### Resources Table

| REAGENT or RESOURCE | SOURCE | IDENTIFIER |
| --- | --- | --- |
| **Experimental Models: Cell Lines** |  |  |
| HEK293A | Thermo Fisher Scientific | Cat# R70507; RRID:CVCL_6910 |
| *HEK293T* |  |  |
| **Recombinant DNA** |  |  |
| pLenti-DsRed-IRES-eGFP | Addgene | Cat# 92194; RRID:Addgene_92194 |
| Human Kinome CRISPR Knockout Pooled Library (Brunello) | Addgene | Cat# 1000000083; RRID:Addgene_1000000083* |
| psPAX2 (lentiviral packaging plasmid) | Addgene | Cat# 12260; RRID:Addgene_12260 |
| pMD2.G (envelope plasmid) | Addgene | Cat# 12259; RRID:Addgene_12259 |
| **Chemicals, Peptides & Recombinant Proteins** |  |  |
| *Bovine calf serum* | *VWR Life Science Seradigm* | *TBD – catalog # not given in manuscript* |
| MEM Vitamins (100X) | VWR | Cat# 45000-702; RRID: N/A |
| PolyJet transfection reagent | FroggaBio | Cat# SL100688; RRID: N/A |
| Polyethylenimine (PEI) | Polysciences | Cat# 23966; RRID: N/A |
| Polybrene | MilliporeSigma | Cat# H9268-5G; RRID: N/A |
| Crystal violet | Bio Basic | Cat# CB0331; RRID: N/A |
| PBS | Wisent | Cat# 311-010-CL; RRID: N/A |
| Paraformaldehyde (PFA) | Bio Basic | Cat# PB0684; RRID: N/A |
| EDTA | Thermo Fisher Scientific | Cat# BP118-500; RRID: N/A |
| Tris | Thermo Fisher Scientific | Cat# BP152-5; RRID: N/A |
| BSA | Bio Basic | Cat# D0024; RRID: N/A |
| SDS | Bio Basic | Cat# SB0485; RRID: N/A |
| Proteinase K | New England Biolabs | Cat# P8107S; RRID: N/A |
| RNase A | Bio Basic | Cat# RB0473; RRID: N/A |
| Ammonium acetate | Bio Basic | Cat# AR0032; RRID: N/A |
| Isopropanol | Thermo Fisher Scientific | Cat# 383910025; RRID: N/A |
| Ethanol | Greenfield Global | Cat# P016EAAN; RRID: N/A |
| *G418 (Geneticin)* | *Not specified in manuscript* | *TBD – please provide vendor/catalog #* |
| *Puromycin* | *Not specified in manuscript* | *TBD – please provide vendor/catalog #* |
| **Critical Commercial Kits & Consumables** |  |  |
| 2X Q5 Master Mix | New England Biolabs | Cat# M0494L; RRID: N/A |
| AMPure XP Bead-Based Reagent | Beckman Coulter | Cat# A63881; RRID: N/A |
| Nextera XT Index Kit | Illumina | Cat# FC-131-1001; RRID: N/A |
| PEG-based Virus Precipitation Kit | Benchmark Bioscience | Cat# P-250; RRID: N/A |

## Methods

### 1. Summary of screen workflow

A CRISPR screen workflow is summarized in the steps below and is illustrated in Figure 1. Preparatory work, optimization and technical details for each step are expanded upon in the sections indicated

**Figure 1.**
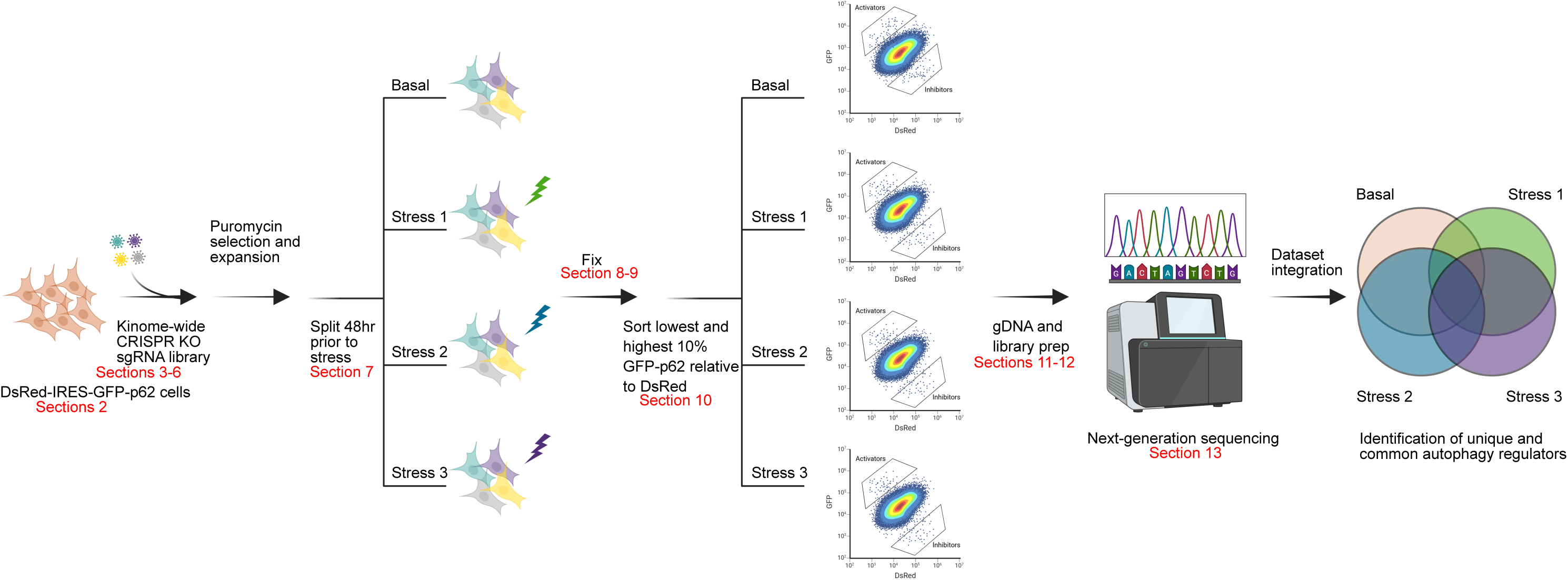
Workflow of kinome-wide CRISPR screen for the identification of selective autophagy regulators

**Day 0-** Reporter cells are plated in a quantity that ensures they reached the desired cell number and confluency at the time of transduction. See *Section 2* for instructions on generating reporter cells.

**Day 1**- **Viral transduction**. Reporter cells are transduced with a pooled library of lentiviruses encoding CRISPR-Cas9 editors in the presence of polybrene (10 µg/mL). See *Sections 3, 4, and 6* for viral production and titering instructions.

**Day 2-3-** Transduced cells will likely require passaging to avoid confluence or cell cycle arrest. Cells should be split such that they adhere and grow to approximately 50% confluence before drug selection in the following step.

**Day 3- Drug selection of transduced cells.** Positive selection for infected cells should be performed by antibiotic resistance. Recommendations regarding the use of kill curves and frequency of media changes for reporter line production is found in *Section 2*

**Day 3-Day 16-** Transduced reporter cells are cultured in the selection media until the genetic disruptions are reflected at the proteomic level. The length of time should be determined empirically using positive and negative controls in your reporter line. See *Section* 2 for determining knockout efficiency in the reporter line.

**Day 16- Plate reporter cells for treatment**- The transduced cells are split and plated for experiment. The plating must achieve a desired cell count and confluency to maintain library representation (*See section 7*).

**Day 18- Experimental treatment**- Stress is applied using experimentally determined timepoints, which yield consistent reporter response (see *Section 2 for more details*).

**Day 18- Fixation-** Following the treatments, aspirate the medium, wash the cells once with PBS, and fix with 2% PFA. The sensitivity of your cells and reporter fixation conditions should be empirically determined (see *Section 8 for more details*).

**Day 18-Cell Harvesting-** Aspirate PFA and Tris-HCl and wash cells once with PBS. Add ice-cold flow buffer, detach cells using a cell scraper and filter cells through a cell strainer (70 µm; Falcon, cat. no. CA21008-952) (see *Section 8 for more details*).

**Potential pause point-** Fixed cell suspensions can be stored at 4°C. Storing fixed cell suspensions overnight at 4oC protected from light retained sufficient reporter fluorescence for sorting.

**Post fixation-Sorting-** FACS procedure will depend on equipment and software availability, but key parameters such as speed and length should be defined prior to screening (see *Sections 9 and 10*). Gating stringency will depend on study specific parameters and may include both high and low populations (see *Section 10*). During sorting, the total number of sorted cells must be recorded to verify the representation of each sample.

**Post fixation - Cell Collection-** Sorted and unsorted populations are pelleted by centrifugation (4000 rpm for 10 min at 4°C). The supernatant is aspirated and cell pellets are stored at −80 °C.

**Repeat-** These steps should be repeated for all biological replicates before proceeding to DNA extraction. Some protocols recommend a minimum of two biological repeats per condition, but three or more are considered best practice for appropriate statistical analysis and reproducibility^30,31^.

**Post fixation-DNA extraction-** Genomic DNA from fixed cell pellets is extracted according to the protocol in *Section 11*.

**Post fixation-PCR amplification and barcoding-** To prepare the samples for next-generation sequencing, guide regions need to be amplified and barcoded. PCR reactions and purifications are conducted according to instructions provided in *Section 12*. PCR products are stored at −20°C or −80°C until sequencing (see *Section 13*).

### 2. Engineering a versatile reporter cell line for high throughput screening

#### Reporter selection

Several factors should be considered when designing an optimal screen reporter. Firstly, an optimal reporter should exhibit stable and uniform expression throughout the cell population (i.e., a monoclonal cell line). A monoclonal reporter line, rather than polyclonal, minimizes reporter variability from population drift, thereby improving reproducibility. The level of reporter expression must also be considered. The reporter needs to be expressed in high enough quantities to have sufficient signal for detection by flow cytometry, but not to highly expressed such that it no longer responds to stress conditions similarly to the endogenous protein. Different promotors driving reporter expression provide distinct characteristics (see Table 2). Endogenous tagging of a reporter gene is also an option. However, for autophagy proteins, we have found endogenous tagging often yields insufficient signal for flow cytometry. Additional considerations include determining the sensitivity of the reporter to cell fixation (*Section 8*) and its impact on cell growth and viability.

**Table 2.**
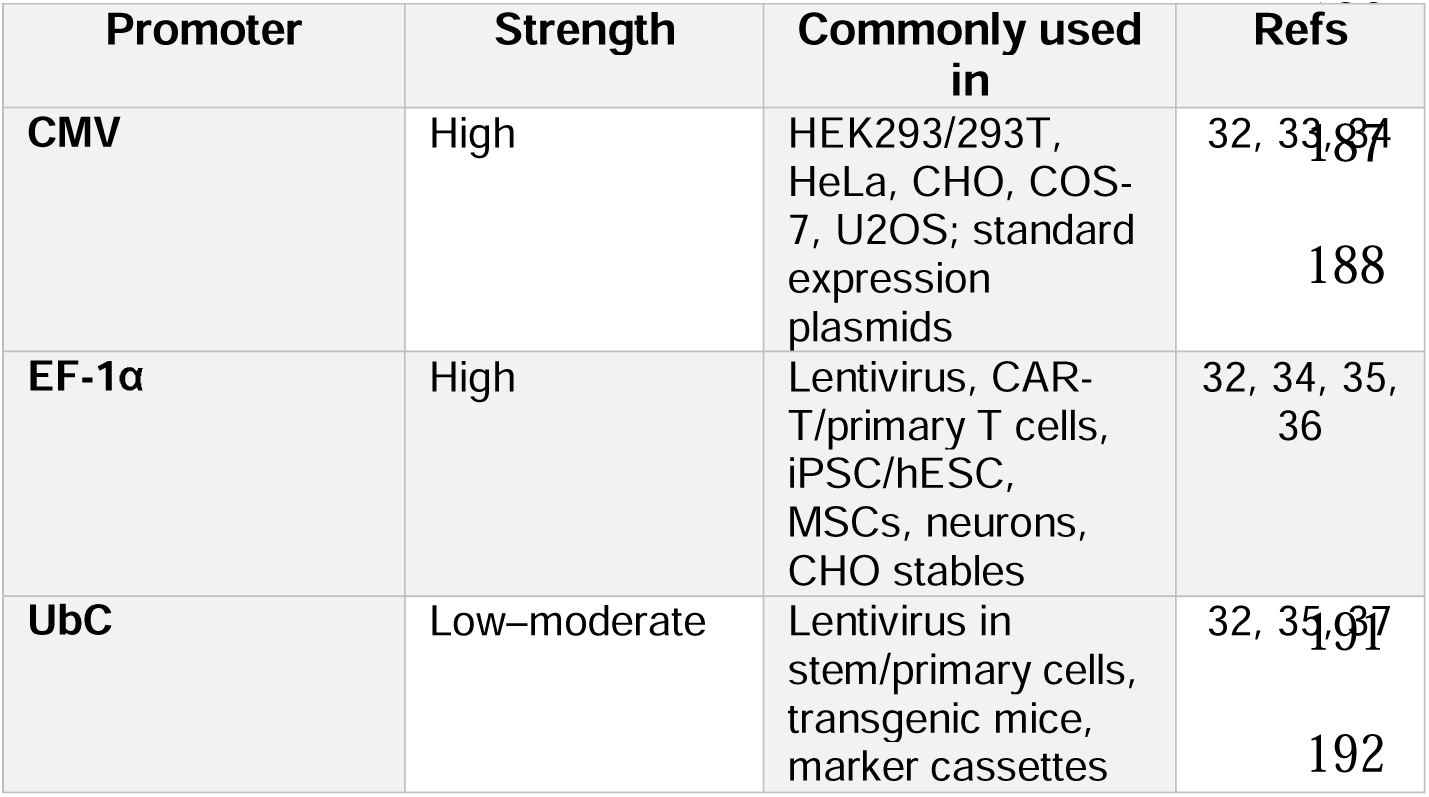
Comparison of promotors for reporter expression.

#### Design of a DsRed-IRES-GFP-p62 autophagy reporter

In our recent study we aimed to uncover signaling pathways governing selective autophagy. We selected sequestosome 1 (p62) as our readout, because of its role in many forms of selective autophagy and the well-established inverse correlation between p62 protein stability and autophagy flux^19,38,39^. Mechanistically, p62 acts as a cargo adaptor targeting ubiquitinated substrates to the lumen of the forming autophagosome and as such, it is degraded along with the autophagy cargo^17,19^. To enable internally normalized and high throughput quantification of p62 dynamics, we engineered a dual fluorescence reporter, DsRed IRES GFP p62 (Figure. 2).

**Figure 2.**
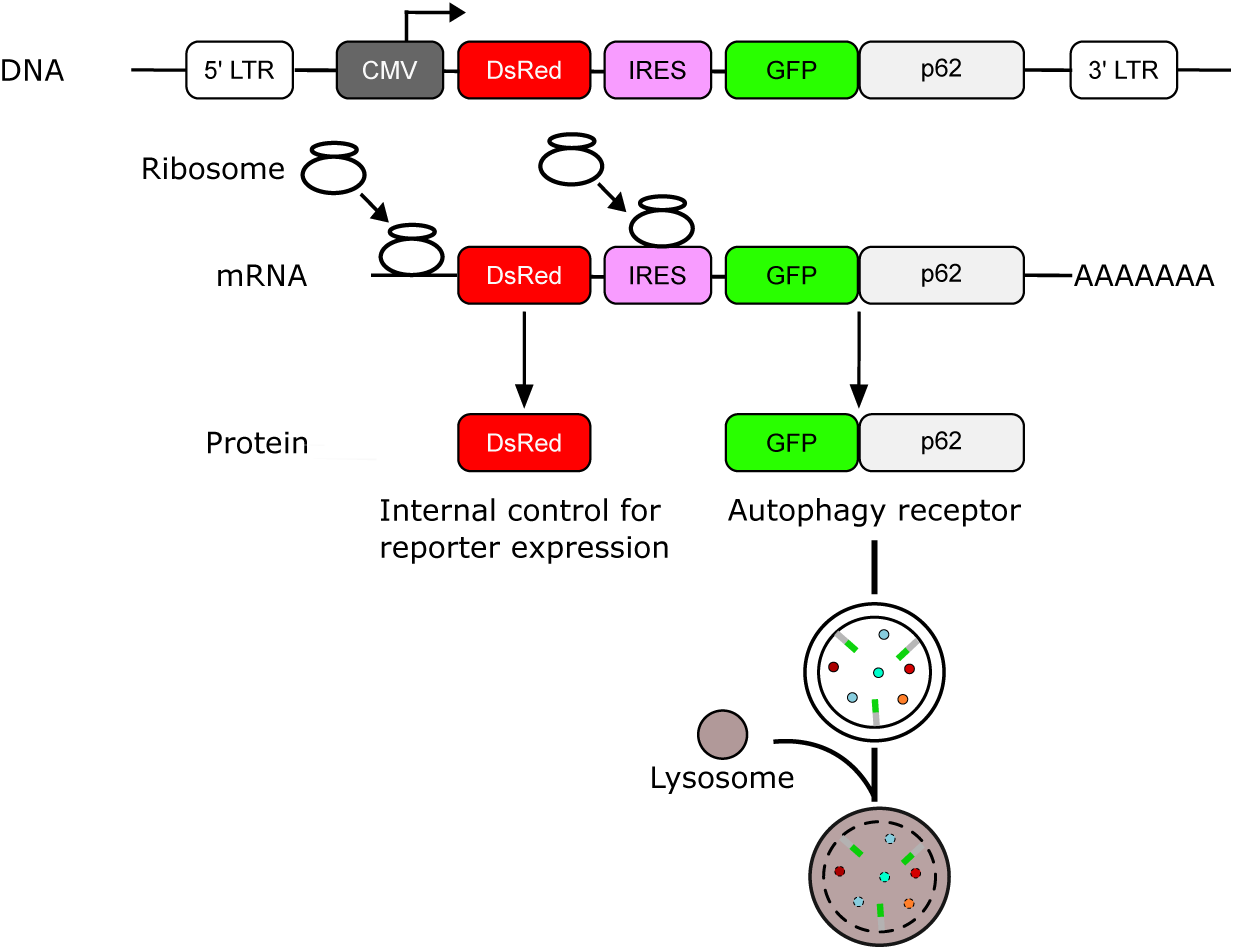
Reporter diagram. The schematic depicts the autophagic flux reporter consisting of DsRed and GFP-tagged p62. p62 is selectively integrated into and degraded alongside the autophagosomal membrane. Thus, the expression level of GFP-p62 correlates inversely with autophagic flux.

When autophagy is activated under stress conditions, p62 levels decrease due to lysosomal degradation, which quenches the GFP signal. Meanwhile, DsRed expression remains stable and unaffected by autophagic flux, providing an internal reference for p62 expression. The result is that any shift in GFP p62 signal, measured by flow cytometry or western blot, is more likely to reflect true changes in p62 degradation rather than transcriptional or translational changes in reporter expression.

#### Production of lentiviruses carrying DsRed-IRES-GFP-p62

The DsRed-IRES-GFP-p62 was introduced into our cells using lentivirus using the protocol described below. However, reporter integration using the TREX system or piggy bac system^40^ are also viable options.

1. 3 µg of plasmid containing DsRed-IRES-GFP-p62, 1 µg of psPAX2, and 0.25 µg of pMD2.G was used to transfect one 10 cm plate of HEK293T cells at 35% confluence (3-4 million cells) using 17 µL PEI (1 µg/µL). Media was changed 5 hours post transfection to avoid cytostatic effects of PEI.
2. Media was collected four times, 12 hours apart, starting at 36 hours post transfection. Media was stored at 4°C until all four fractions were collected.
3. Viral supernatant was filtered through 0.45 µm Nalgene™ Rapid-Flow™ Sterile Disposable Filter Units (Thermo Fisher Scientific#09-740-63A). If a significant number of detached cells are present in the collected media, centrifugation at 300 × g for 5 min should be employed to reduce filter clogging. Cleared supernatants can be used directly for infection or concentrated using a PEG-based Virus Precipitation Kit (#P-250 Benchmark Bioscience) to reduce to 1/100 of the original volume. Using concentrated virus preparations can be advantageous or even necessary when working with cell lines that are difficult to transduce. By concentrating the virus, the target transduction rate is more readily achieved, facilitating the establishment of stable cell lines and ensuring sufficient genetic modification for downstream applications. This also allows for the storage of high titer viral preps in the freezer, improving the reproducibility of replicate screens conducted at later times.
4. The viruses were stored at −80°C.

#### Generation of HEK293A expressing DsRed-IRES-GFP-p62

1. We chose HEK293A cells as the preferred background for developing our reporter cell line. HEK293T cells were not used because they are inherently resistant to G418 and have inferior morphology for microscopic analysis. If creating stable cell lines for the first time in a cell background, perform a kill curve with the non-transduced line and ensure the selection method will not impair selection in downstream steps (i.e. drug resistance of CRISPR library).
2. Virus transduction of HEK293A cells was performed in 6-well format at 40% confluence.
3. Virus was added to HEK293A cells in the presence of polybrene (10 µg/mL). The viral titer can be increased for difficult to infect cells and polybrene concentration can be reduced for sensitive cells (typically used from 1 to 10 µg/mL). The following day, cells were transferred to 10-cm plates.
4. 48 hours post transduction, transduced cells and a negative control uninfected plate were treated with antibiotics. Specifically, cells were cultured in DMEM containing G418 (1 mg/mL) for 7 days, or until all control cells die. The culture media were replaced every day with G418-containing media. If more than 5% of the original population of cells have died since the last media change, another media change is needed. The presence of too many dead cells or apoptotic debris can impact growth and viability of transduced cells and should be avoided. Cells should be monitored daily until selection is complete as each cell line and antibiotic may impact the timeframe of selection.
5. After recovering from drug selection, the reporter cell line underwent functional validation to confirm its continued responsiveness within the signaling pathways of interest.

#### Validation of reporter line

Validating the specificity and biological relevance of a reporter cell line is a critical step before it is employed in pooled genetic screens or functional assays. This ensures that changes observed in the reporter signal reflect the intended cellular process rather than unrelated pathways or experimental artifacts. Specificity is tested by perturbing well-characterized positive and negative regulators of the pathway of interest and confirming that the reporter responds as expected using orthogonal detection methods such as western blot or flow cytometry. For example, to validate the specificity of our DsRed-GFP-p62 reporter, we knocked out an established activator of starvation-induced p62 turnover, unc-51 like autophagy activating kinase 1 (ULK1)^24,41–44^.

#### Monitoring reporter cell responses to perturbations in cellular homeostasis

After confirming the specificity of reporter expression, it is essential to optimize the stress conditions relevant to the study^45–53^. Selecting the appropriate time point is a critical part of the screening process; ideally, this time point should capture the strongest response as early as possible to minimize potential secondary effects. While longer time points might yield a greater number of candidate hits, they risk including indirect or downstream consequences unrelated to the primary pathway regulation. In contrast, acute time points are more likely to reflect direct involvement in pathway modulation.

#### Validation of DsRed-IRES-GFP-p62 reporter line

In the context of our study, we began by evaluating the expression and responsiveness of a polyclonal DsRed-GFP-p62 reporter population by western blot. Amino acid starvation was used to induce autophagic degradation of p62^45–47^. We found that the polyclonal cells exhibited highly elevated levels of DsRed-GFP-tagged p62 compared to the endogenous p62 and showed no detectable responses to amino acid starvation. This shows how high overexpression changes the stoichiometric ratio of p62 to its regulators and blocks our ability to detect p62 turnover (Figure 3A). To solve this, we sorted the DsRed-GFP-p62 reporter line into monoclonal populations and evaluated them western blot. We found a monoclonal reporter clone that showed consistent and expected responses of p62 flux to amino acid starvation (Figure. 3B).

**Figure 3.**
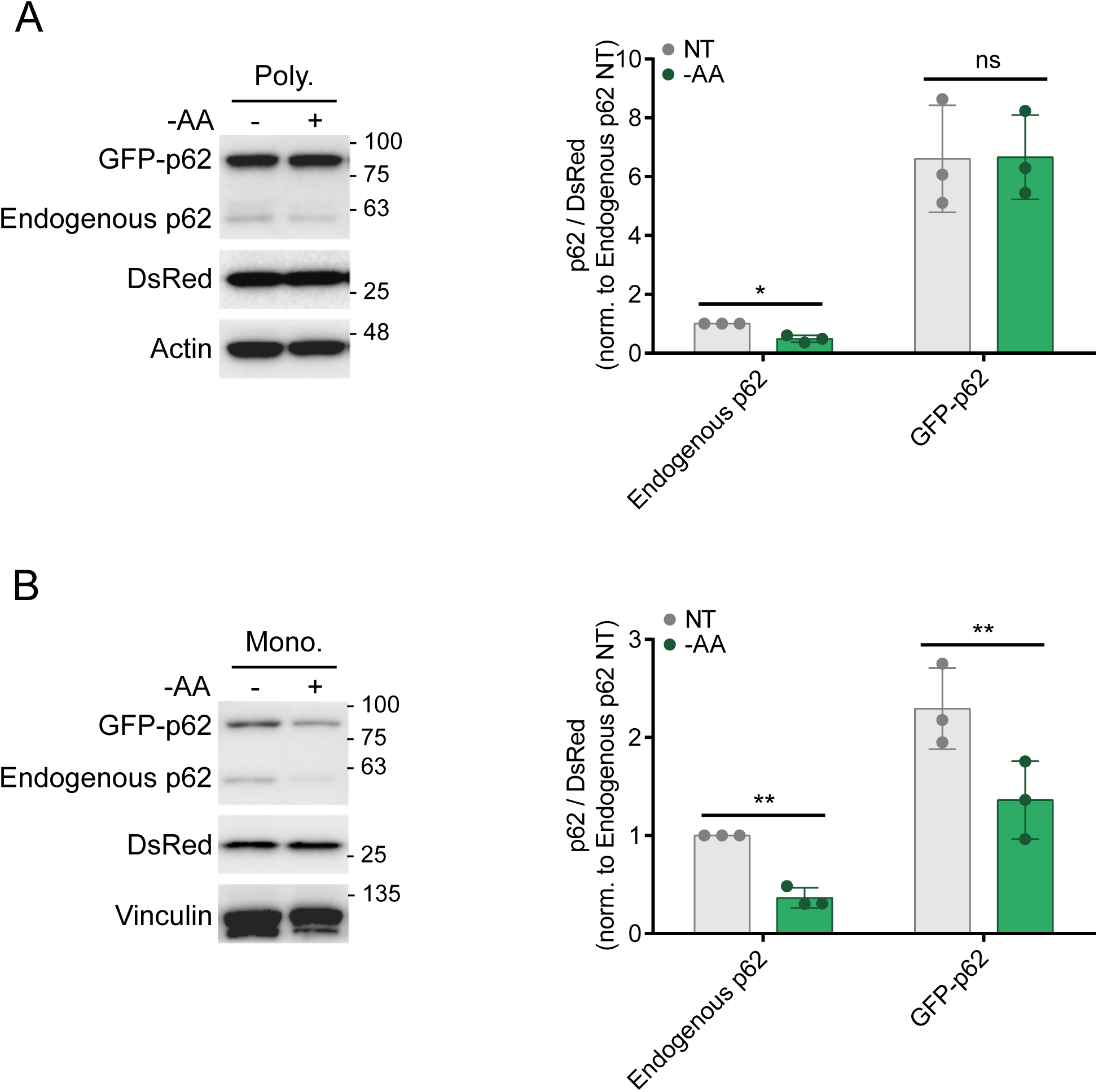
Examination of autophagy flux through examination of p62. **(A), (B)** Polyclonal (**A**) and optimal monoclonal (**B**) populations expressing DsRed and GFP-p62 were incubated with amino acid-free media for 3 hours. Whole-cell lysates were immunoblotted using the antibodies indicated.

### 3. Lentiviral preparation for delivery of the genetic perturbation library

When working with pooled libraries, whether targeting specific gene families or genome wide collections, it is essential to begin by amplifying the library according to the provider’s recommended protocol to ensure high quality representation. Adequate library coverage, typically on the order of at least 1000 fold, should be confirmed prior to virus production, as insufficient representation can compromise the robustness and reproducibility of downstream experiments. In our workflow, the selected human kinome CRISPR knockout pooled library was obtained from Addgene and amplified following the supplied protocol^54^. Library representation was then assessed using a two step PCR method (described below), and the resulting amplicons were submitted for next generation sequencing (NGS). NGS datasets are analyzed using various computational tools; in this study, we used the CRISPRBetaBinomial (CB2) method via the CRISPRCloud2 web platform^55^. CB2 is a web based analysis tool that provides key quality metrics, including total read counts, mapped reads, mapping efficiency, and fold coverage, offering a clear assessment of library quality and representation before moving forward in the screening pipeline. The procedure for producing lentiviruses carrying the kinome library is described below.

1. For a library size of roughly 3,200 guides, we infected 6-8 15 cm plates of HEK293T cells at approximately 35% confluency (7-8 million cells). The amount of virus producing cells depends on the number of viruses required in *Section 6*.
2. Next day, lentiviral vectors expressing the kinome library and packaging vectors (psPAX2 and pMD2.G) were co-transfected using PolyJet reagent into HEK293T cells in a 4:3:1 molar ratio, respectively. 3 μL of PolyJet was used for 1 μg of DNA. Media were changed 5 hours post transfection.
3. Media were collected four times 12 hours apart starting at 36 hours post transfection.
4. Viral supernatant was filtered through 0.45 µm Nalgene™ Rapid-Flow™ Sterile Disposable Filter Units (Thermo Fisher Scientific#09-740-63A). Cleared supernatants were concentrated using a PEG-based Virus Precipitation Kit to 1/100 of the original volume.
5. The viruses were stored at −80°C.

### 4. Quantification of lentiviral particles harboring the screening library

Following lentivirus production, the viruses should be titrated in the reporter cell line. This step is important as it measures the number of infectious viral particles per unit volume of the viral stock and is subsequently used for Multiplicity of Infection (MOI) calculation.

1. Approximately 10-15% confluence (0.8×10^5^) of reporter cells were seeded to each well of a 12-well plate in complete DMEM media containing 10% bovine calf serum without antibiotics and the cells were incubated overnight at 37°C, 5% CO_2_.
2. On the next day, lentiviruses were thawed on ice and resuspended gently. 5-fold serial dilutions of viral stock were performed in a 96-well plate.
3. Culture medium from each well was removed. The reporter cells were replenished with 1 mL fresh media in the presence of polybrene (10 µg/mL).
4. A single viral dilution was gently added to each well of a 12-well plate. The plate was mixed gently from one side to another side.
5. Media were changed to regular complete DMEM 24 hours post transduction. The cells were incubated at 37°C with 5% CO_2_ for another 24 hours.
6. Next, the transduced cells were incubated with puromycin (1 µg/mL) for 3 days. The culture media were replaced every day with puromycin-containing media if more than 10% of cells died.
7. The cells were incubated with regular complete DMEM for another 4-6 days and observed every day to monitor the death of cells that were sensitive to puromycin.
8. The transduced cells were fixed with 4% PFA for 10 min at room temperature.
9. Following one wash with 1x PBS, the cells were stained with 0.1% crystal violet solution at room temperature for 20 minutes. After removal of crystal violet solution, the samples were washed three times with 1x PBS.
10. The blue-stained colonies were counted using a microscope at a magnification of 40×.
11. The lentiviral titer was calculated using the formula below

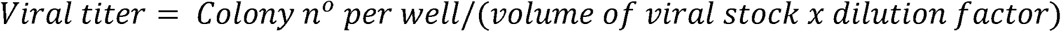

Viral titer (transduction unit per mL or TU/mL): the viral particle count added per cell during infection.
Colony n° per well: the number of colonies in one well.
Dilution factor: a measure of how much viral solution has been diluted.

### 5. Establishing the minimum number of cells required for transduction with lentiviruses containing the library of interest

Ensuring an adequate number of target cells for transduction with viruses carrying the library of choice is essential to maintain robust library representation and reliable screen outcomes. For pooled genetic screens, such as those using CRISPR-Cas9 knockout libraries, it is recommended to transduce a sufficient cell population to achieve at least 500-1000× coverage for each library element^9,56,57^. The minimum number of reporter cells required for each condition should be calculated based on library size and desired coverage, as outlined below. Note that a multiplicity of infection (MOI) of 0.3 is routinely used to ensure that the majority of reporter cells receive a single copy of guide RNA^9,30,57^.

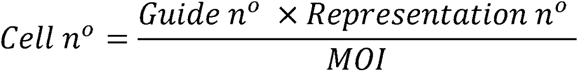

Cell n°: the minimum number of reporter cells to be infected.
Guide n°: the total number of guides in the library, including non-targeting controls.
Representation n°: the expected representation (usually 1000x).
MOI: multiplicity of infection.

For example, our library size of 3200 guides and 1000x representation requires at least 10.7 million cells (50-60% of a 15-cm plate) for viral transduction per treatment condition.

### 6. Determining the required viral volume for reporter cell transduction

The number of lentiviral particles required to transduce the reporter cell line is determined based on the desired MOI, the total number of target cells, and the viral titer, according to the formula:

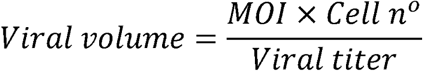

Viral volume (mL): the number of viruses added to the reporter cells.
MOI: Multiplicity of infection
Cell n°: the total number of cells measured immediately prior to viral transduction, which should meet or exceed the minimum value calculated in Section□5.
Viral titer (TU/ml): determined in Section 4.

### 7. Calculating the number of transduced cells required for sorting

The number of pre-sort cells needed for each condition depends on library size, desired guide representation, gating, and cell type and confluency. The following calculation provides the number of culture plates needed to maintain a 500-1000x guide representation post sort.

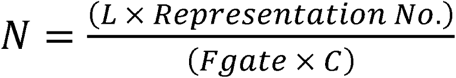

N=number of plates required
Representation No.=representation
L=Library size (guide number)
F_gate_=fraction of events falling in the sort gate
C=Number of cells per plate

For example, the Brunello kinome library consists of 3200 gRNAs. We would like to maintain 1000 fold representation. We gated the top and bottom 10% of cells for GFP:DsRed signal, making our gating fraction 0.1. HEK293A cells at 80% confluency yield approximately 2×10^7^ cells per 15cm plate.

The calculation would be as follows:

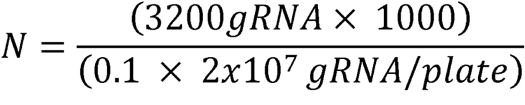

*N* = 1.6 plates

Therefore, two 15cm plates of cells are necessary for each condition. Two plates also provide a buffer for loss of cells during treatment and fixation.

### 8. Optimization of crosslinking conditions

Crosslinking or fixation can cause changes to fluorescent signals. In our study, we utilized paraformaldehyde (PFA), a commonly used crosslinking reagent, to fix samples. Additionally, we used Tris to quench formaldehyde reactivity to prevent excess crosslinking. We investigated various fixation conditions using PFA in the presence or absence of Tris and examined fluorescence of fixed samples using flow cytometry (Figure. 4). We observed that higher concentrations of PFA and longer treatment times resulted in deteriorated fluorescence (Figure 4). Moreover, Tris addition helped prevent quenching of the GFP signal (Figure. 4). The optimal conditions identified for our screen are described below, but may vary with fluorophore and expression levels.

**Figure 4.**
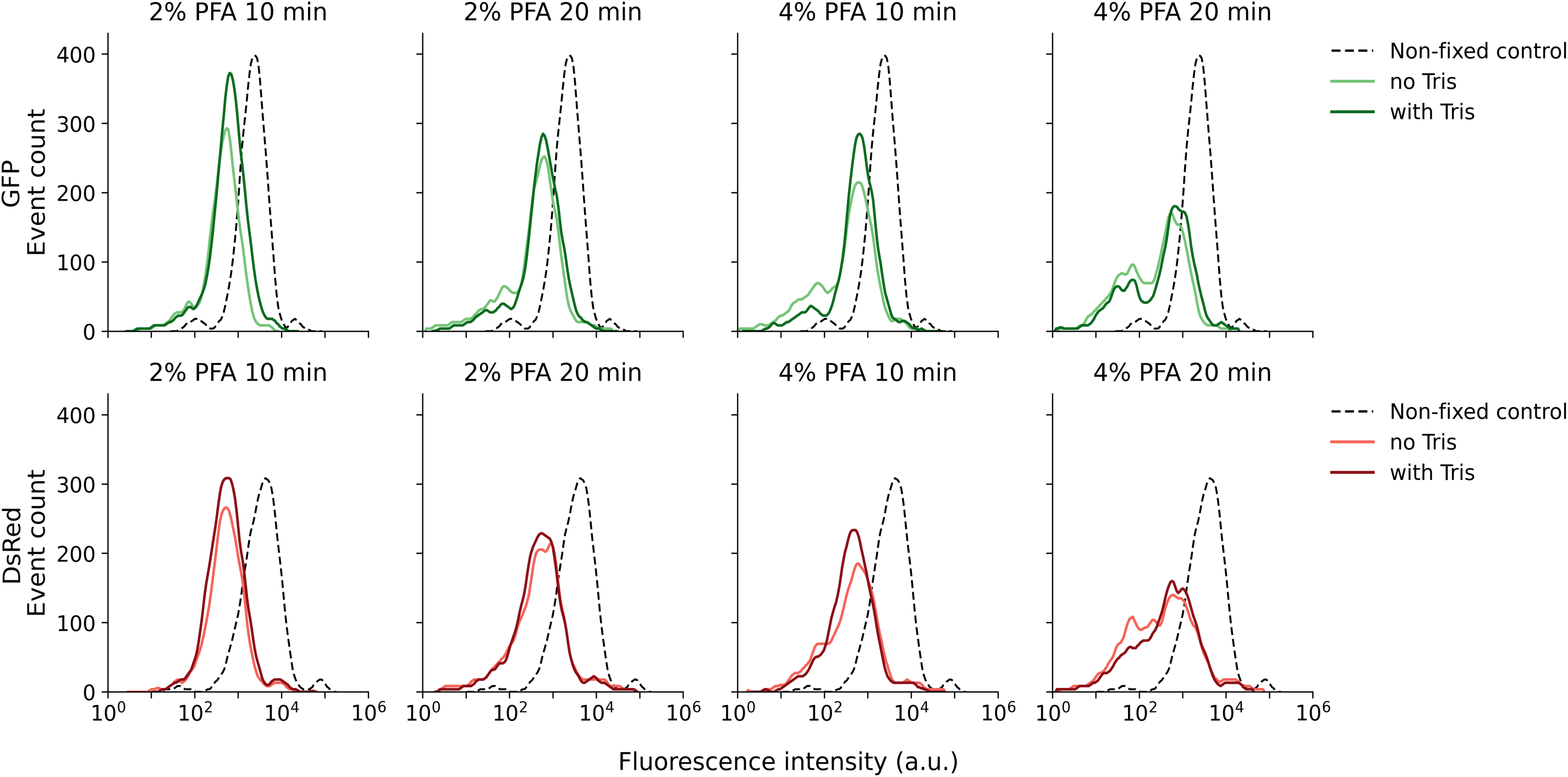
Crosslinking/fixation optimization. The reporter line underwent fixation with either 2% or 4% PFA for durations of 10 or 20 minutes, with or without the addition of Tris, at room temperature. Subsequently, the fixed cells were subjected to flow cytometry analysis.

In our screen, we used 2% PFA for 10 min at room temperature. Crosslinking was stopped with the addition of Tris-HCl (pH 8, final concentration 1 M) for 15 min

1. Sample media were aspirated. The cells were washed once with PBS.
2. The cells were incubated with 2% PFA for 10 minutes at room temperature, followed by 15 min incubation with 3 M Tris (pH 8, direct addition to 2% PFA to create a final concentration of 1 M).
3. Tris and PFA solution were removed. The samples were washed once with PBS.
4. After PBS removal, the cells were collected using scrapers, stored in ice-cold flow buffer (1% BSA and 2 mM EDTA in PBS), and filtered using 70 µm cell strainers (Falcon, cat. no. CA21008-952).
5. These fixed cells were stored at 4°C until analysis by flow cytometry.

With this approach, we observed sufficient fluorescent signals in the fixed samples with minimal reduction in signal intensity in response to PFA fixation.

### 9. Benefits of cell fixation for extended sort times

We then investigated how the duration of wait times prior to sorting affected autophagy flux. Our data revealed that p62 levels declined even at the earliest measured timepoint, consistent with rapid autophagy initiation after the samples were placed on ice (Figure. 5A). Notably, the response kinetics differed between selective autophagy receptors: RETREG1/FAM134B (an ER-phagy receptor)^48^ and PMP70 (a peroxisomal membrane marker)^51,52^ exhibited distinct temporal activation patterns (Figure 5A). This highlights the importance of considering variable pathway activation rates when comparing different types of selective autophagy, since apparent discrepancies may arise from distinct temporal responses to prolonged wait times rather than fundamental biological differences. Additionally, cells may experience stress upon detachment and exposure to FACS buffer, which induces autophagic degradation of p62, thereby diminishing the effectiveness of the screen read-out.

**Figure 5:**
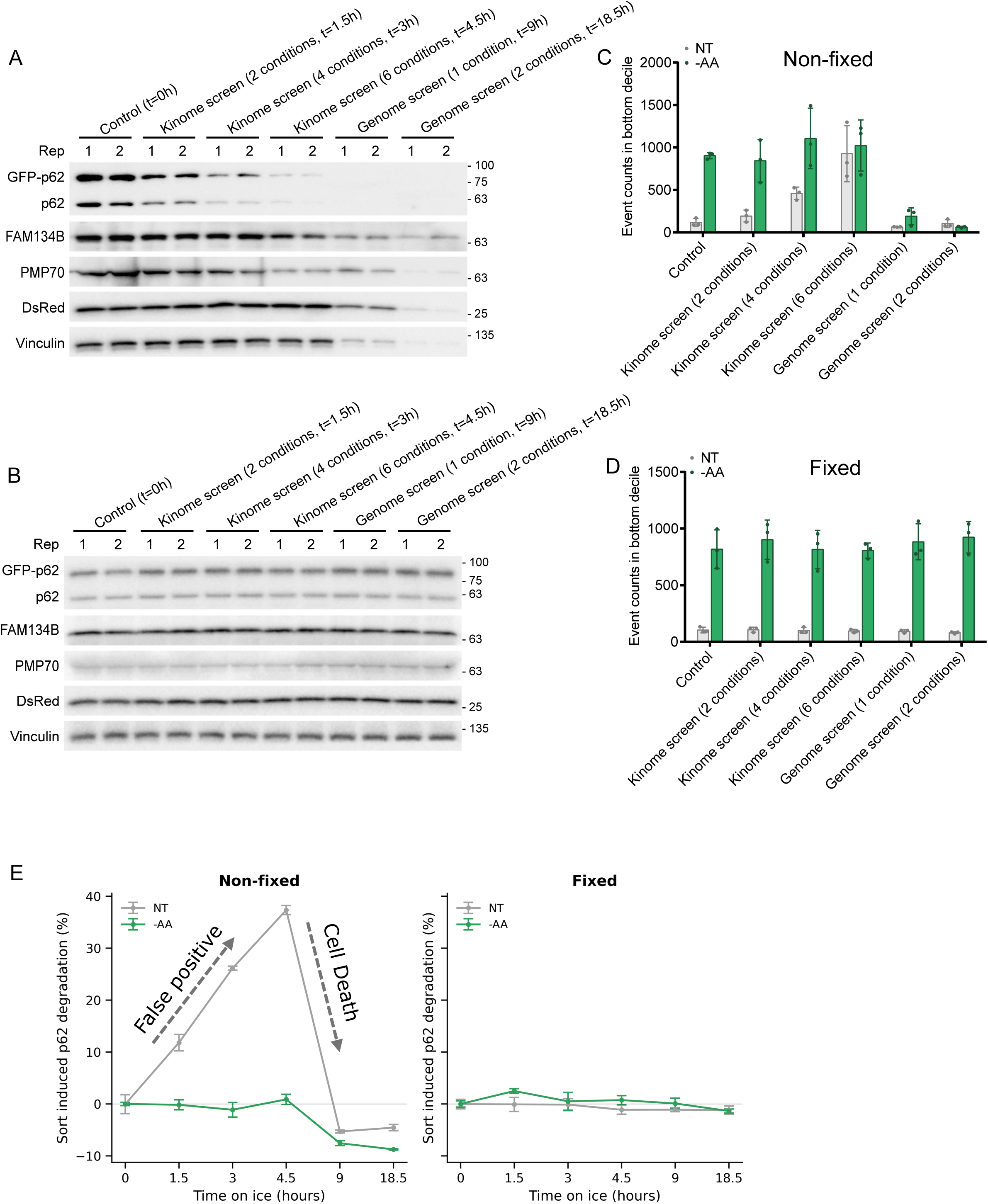
Comparison of non-fixation and fixation procedures. (**A**), (**C**) Reporter cells were either analyzed immediately or held on ice for durations equivalent to the corresponding screen protocol before being assessed by western blot (**A**) or flow cytometry (**C**). (**B**), (**D**) Reporter cells were fixed prior to analysis. The fixed samples were then either analyzed immediately or stored on ice for the same protocol-matching intervals before analysis by western blot (**B**) or flow cytometry **(D)**. **(E)** Quantification of sort-induced p62 degradation by flow cytometry for fixed and non-fixed cells during incubation on ice.

To resolve these stress-induced artifacts, we incorporated a sample fixation step into our workflow. We found that crosslinking effectively preserved autophagy markers such as p62, RETREG1, and PMP70, preventing signal loss during prolonged ice incubation (Figure 5B). Using flow cytometry, we evaluated the impact of extended sample handling (without or with fixation) and included starved samples as controls (Figure 5C-D). Using the bottom 10% of cells from starved, immediately processed samples as a reference gate, we assessed whether stress from prolonged wait times shifted cells into this low-signal population. Consistent with our western blot data, event counts in the bottom decile increased over time for non-fixed samples under basal conditions compared to fixed counterparts, suggesting that extended wait times may elevate false-positive rates and thereby reduce the statistical power of screen analysis (Figure 5C, E). After 4.5 hours on ice, the number of events included in the bottom decile for unfixed basal and starved conditions rapidly declined, consistent with progressive cell death. In contrast, we observed no change in p62 signal in fixed samples after 4.5 hours on ice (Figure 5D-E). Together, these results indicate that fixation preserves signal integrity during long workflows and reduces the contribution of handling artifacts to the screen readout.

### 10. Sorting of targeted populations

Cell sorting by fluorescence-activated cell sorting (FACS) is utilized to separate subpopulations exhibiting changes in the targeted pathway caused by gene disruption. In our screen from Losier et al., we used sorted cells based on the ratio GFP:DsRed fluorescence. As mentioned in Section 2, we used a dual fluorescent reporter for our kinome-wide screen. Specifically, p62 was tagged with DsRed and GFP. DsRed acts as an internal control for p62 expression. Therefore, cells with a lower GFP:DsRed ratio are experiencing higher autophagic flux, while cells with a higher GFP:DsRed ratio are showing less autophagic flux. The sorting gates were designed to capture the top and bottom 10% of cells according to their GFP:DsRed ratio to identify autophagy activators and inhibitors. Gating based on GFP:DsRed ratio for an untreated and amino acid starved population of cells is shown in Figure 6. Depending on the assay sensitivity and biological context, gating thresholds typically range from 5% up to 20% to isolate meaningful subsets of edited cells.

**Figure 6:**
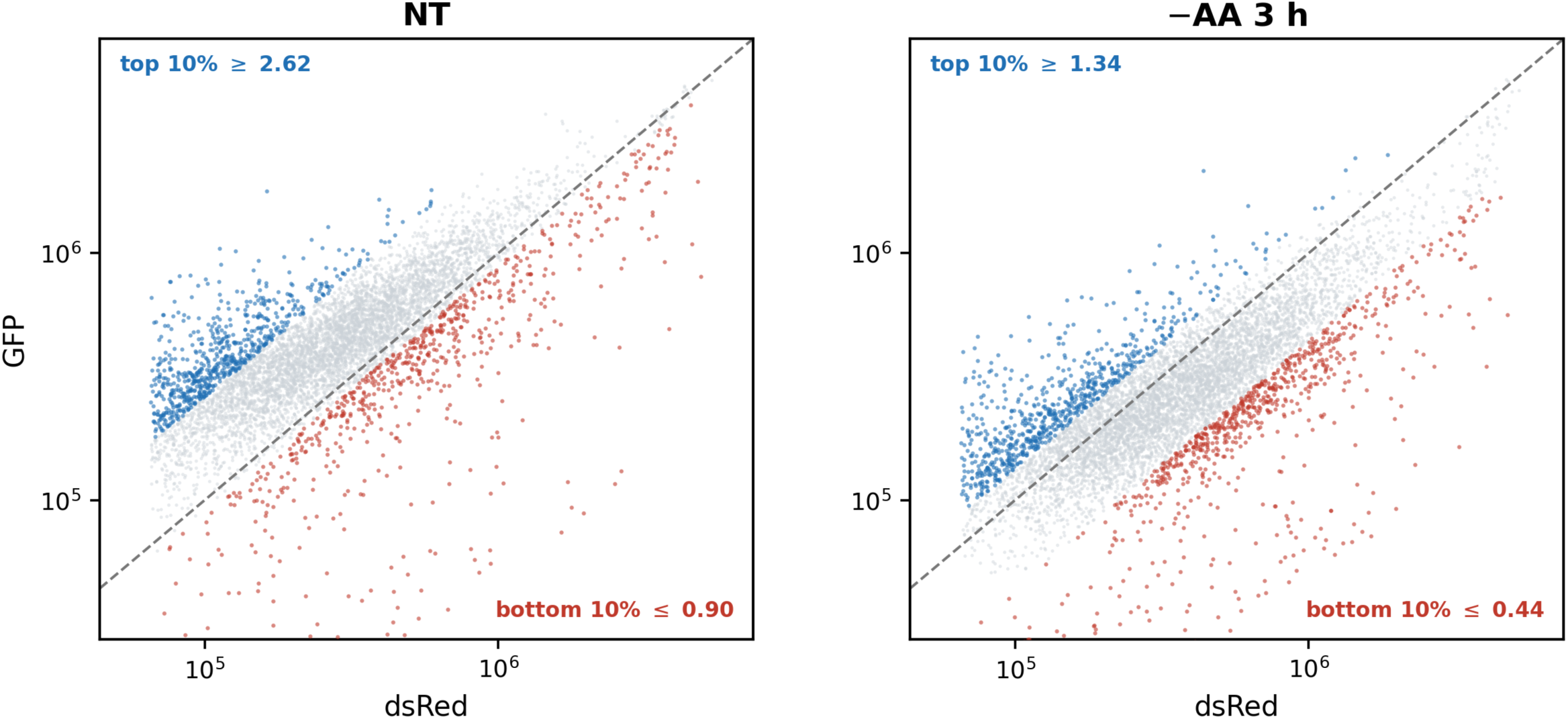
GFP vs DsRed decile gates. FACS gating of the bottom 10% (red) and top 10% (blue) of the GFP:DsRed ratio distribution, calculated for each sample. The left panel is a population of untreated cells; the right panel is amino acid starved cells.

### 11. Genomic DNA extraction from fixed cells

Genomic DNA (gDNA) recovery from fixed cells can be challenging, but efficient recovery is critical to maintaining representation in the screen, therefore a protocol optimized for crosslinked gDNA recovery is used. This protocol allows direct gDNA extraction from up to 2 million fixed cells (further scalable), without a reverse crosslinking step, thereby enhancing reproducibility.

1. In a 15-ml tube, flow buffer was carefully removed.
2. 600 μL of Lysis Buffer (50 mM Tris, 50 mM EDTA, 1% SDS, pH 8) and 3 μL of Proteinase K (20 mg/ml) were added to the cell sample and the mixture was incubated at 55°C overnight.
3. The next day, 1.5 μL of RNase A (20 mg/mL) was added to the lysed sample, which was then inverted 25 times and incubated at 37°C for 30 minutes.
4. The sample tube was cooled on ice for 15 min before addition of 200 μL of pre-chilled 7.5 M ammonium acetate to precipitate proteins.
5. After adding ammonium acetate, the samples were vortexed at high speed for 20 seconds and then centrifuged at 10,000 x g for 10 minutes at 4°C.
6. After the spin, a tight pellet was visible in each tube and the supernatant was carefully decanted into a new tube.
7. 600 μL of 100% isopropanol was added to the tube, which was inverted 50 times and centrifuged at 10,000 x g for 10 minutes. gDNA was visible as a small white pellet in each tube.
8. The supernatant was discarded, 600 μL of freshly prepared 70% ethanol was added. The tube was inverted 10 times, followed by centrifugation at max for 1 minute.
9. The supernatant was discarded by pouring; the tube was briefly spun, and remaining ethanol was removed using a P200 pipette.
10. After air drying for 10-30 minutes, the DNA changed appearance from a milky white pellet to slightly translucent.
11. At this stage, 50 μL of elution buffer of choice was added, the tube was incubated at 65°C for 1 hour and at room temperature overnight to fully resuspend the DNA.
12. The next day, the gDNA samples were vortexed briefly. The gDNA concentrations were examined using a NanoDrop 2000 Spectrophotometer (Thermo Fisher Scientific) and are stored at −20°C for further analysis.

### 12. Sample preparation for next-generation sequencing (NGS)

This step generates sequencing-ready libraries that reflect the abundance of each sgRNA in the screened population. The integrated sgRNA region is first amplified (PCR1) using primers that bind constant vector sequences flanking the variable sgRNA spacer; these primers carry adapter overhangs for a second PCR (PCR2), which adds the sequencing adapters and sample-specific barcodes. This two-step strategy amplifies all integrated guide sequences and produces amplicons compatible with multiplexed next-generation sequencing, allowing sgRNA representation to be quantified across samples (Fig. 7). The following section details our two-step PCR method, adapted from a previously published screen study

**Figure 7.**
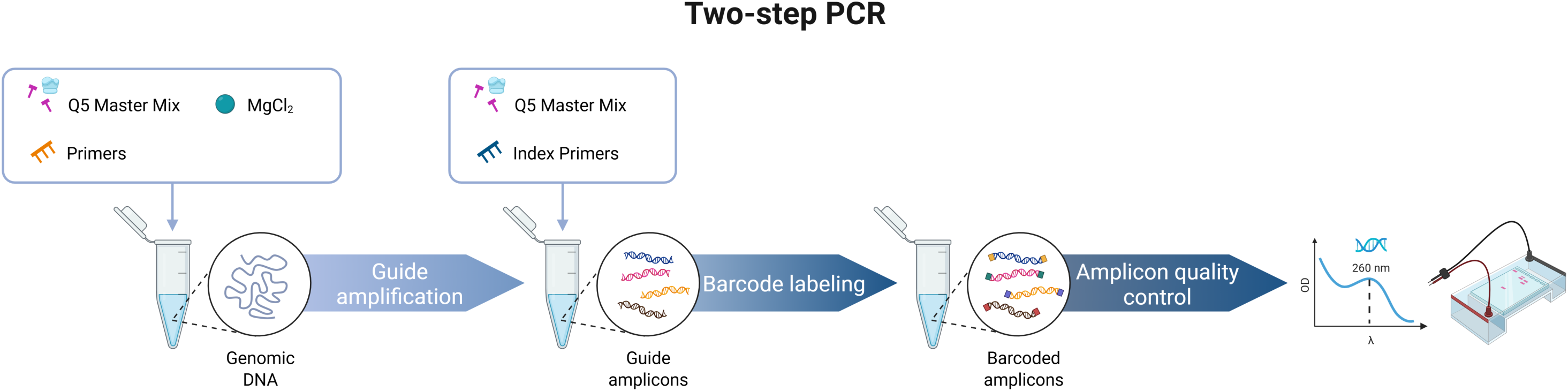
A diagram illustrating the two-step PCR process used to prepare samples for next-generation sequencing.

#### PCR1: amplification of sgRNA target regions from genomic DNA (gDNA)

For PCR1, the total number of reactions and the amount of master mix were calculated once gDNA concentrations and volumes were determined using Excel format. To minimize unexpected variables among samples during PCR, master mix of all reactions were generated by mixing all components except for DNA template and water. For sorted samples, all gDNA was used as a template for PCR1. For bulk or unsorted samples, the amount of gDNA used for multiple PCR1 reactions was determined to guarantee a minimum of 1000x representation was achieved. The amount of gDNA in bulk/unsorted samples can be calculated as follows.

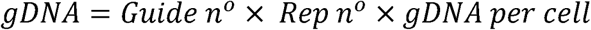

gDNA: the amount of gDNA is used to achieve the expected representation.
Guide n°: the number of guides in the library. Our kinome library has approximately 3200 guides.
Rep n°: the expected representation (usually 1000-1500x).
gDNA per cell: one cell contains roughly 6 pg of gDNA^58^.

PCR1 primers are designed as below.

Forward: TCGTCGGCAGCGTCAGATGTGTATAAGAGACAGggactatcatatgcttaccgt

Reverse: GTCTCGTGGGCTCGGAGATGTGTATAAGAGACAGgagccaattcccactccttt

The capitalized segment of the primer sequence represents a transposase adapter compatible with the Nextera Index Kit utilized in PCR2. The lowercase portion serves as the vector binding sequence and can be adjusted according to the vector of choice. If the vector binding sequence is changed, PCR1 condition should be optimized.

PCR1 mixture and condition are prepared as described below.

|  | Added volume (µL) |
| --- | --- |
| 2x Q5 Master Mix | 50 |
| MgCl<sub>2</sub> (1M) | 0.2 |
| 5' primer (100 µM) | 0.5 |
| 3' primer (100 µM) | 0.5 |
| gDNA (3 µg) | Various |
| H<sub>2</sub>O | Up to 100 |

| PCR1 of gRNA |  |
| --- | --- |
| 98°C | 5 minutes |
| 98°C | 35 seconds |
| 60°C | 30 seconds |
| 72°C | 45 seconds |
| Repeat steps 2-4 for 24 cycles |  |
| 72°C | 10 minutes |
| 4°C | hold |

#### PCR1 cleanup

During this step, AMPure XP beads were employed to isolate the PCR amplicon from genomic DNA, free primers, and primer dimer species. PCR products of each sample from multiple first PCR reactions were pooled and 200 µL is cleaned up for PCR2.

1. The AMPure XP beads were brought to room temperature (minimum 15 minutes) and mixed until homogeneous by vortexing for at least 20 seconds.
2. 0.5X volume (100 µL) of AMPure XP beads (DNA: beads= 1:0.5) was added to respective 1.5 mL Eppendorf tubes. The beads and PCR1 solution were mixed thoroughly and incubated at room temperature without shaking for 15 minutes.
3. Sample tube was placed on a magnetic rack for 2 minutes or until the supernatant has cleared.
4. The supernatant was transferred to a new 1.5mL tube. Beads were discarded.
5. 0.8X volume (160 µL, using original DNA volume) of AMPure XP beads was added. The beads and PCR1 mixture were incubated at room temperature without shaking for 15 minutes.
6. The tube was placed on a magnetic rack for 2 minutes or until the supernatant has cleared.
7. The supernatant was discarded this time, and beads were kept.
8. On a magnetic rack, the beads were washed with freshly prepared 70% ethanol twice according to manufacturer’s instructions.
9. On a magnetic rack, excess ethanol was removed using aspirator and gel loading tips.
10. On a magnetic rack, the beads were air-dried for <15 minutes and checked in 10 minutes to make sure the beads were not over dry.
11. The sample tubes were removed from the magnetic rack. 50-100 µl of ultra-pure water or elution buffer of choice was added.
12. The solution was gently mixed up and down 10 times to make sure the beads were fully resuspended.
13. The mixture was incubated at room temperature for 10-15 minutes.
14. The tube was placed on the magnetic rack for 2 minutes or until the supernatant has cleared.
15. The supernatant was carefully transferred to a new appropriately labeled 1.5 mL tube.
16. 5 µl of PCR1 products were run on a 2% agarose gel to confirm the band size.
17. These samples were analysed by a NanoDrop 2000 Spectrophotometer and were frozen at −20°C before proceeding to PCR2.

#### PCR2: addition of barcodes to PCR1 amplicons using Nextera Index Kit

PCR2 reactions were prepared as below. 62.5 ng of PCR1 corresponds to over 1 billion fold representation.

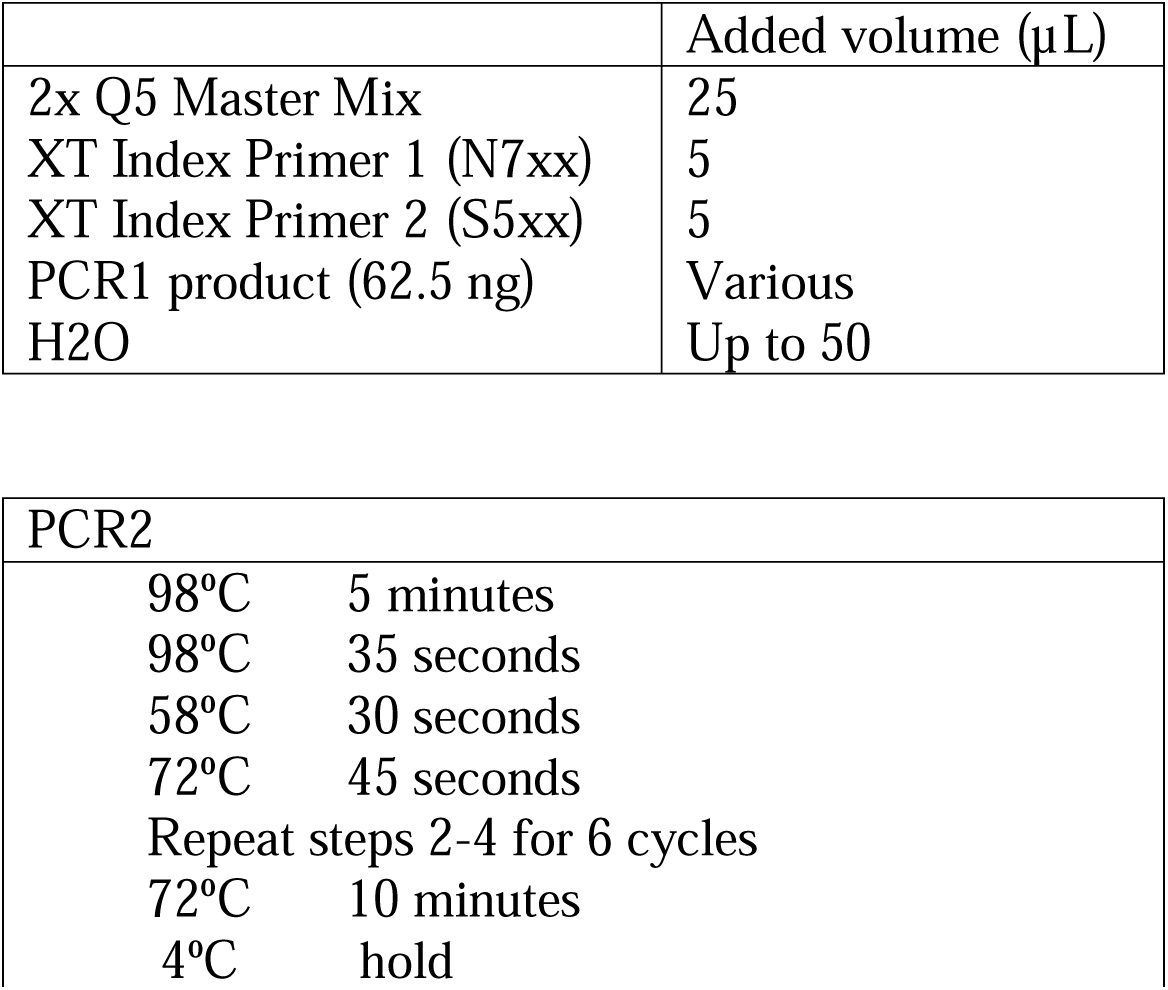

|  | Added volume (µL) |
| --- | --- |
| 2x Q5 Master Mix | 25 |
| XT Index Primer 1 (N7xx) | 5 |
| XT Index Primer 2 (S5xx) | 5 |
| PCR1 product (62.5 ng) | Various |
| H2O | Up to 50 |
| PCR2 |  |
| 98°C | 5 minutes |
| 98°C | 35 seconds |
| 58°C | 30 seconds |
| 72°C | 45 seconds |
| Repeat steps 2-4 for 6 cycles |  |
| 72°C | 10 minutes |
| 4°C | hold |

#### PCR2 cleanup

AMPure XP beads were employed to purify PCR2 products from PCR1 amplicons, primers, and primer dimers. PCR2 cleanup steps are described below.

1. The AMPure XP beads were brought to room temperature (minimum 15 minutes) and mixed until homogeneous by vortexing for at least 20 seconds.
2. 1.6X volume of AMPure XP beads (DNA: beads= 1:1.6) was added to respective 1.5 mL Eppendorf tubes. The beads and PCR2 solution were mixed thoroughly and incubated at room temperature without shaking for 15 minutes.
3. Sample tube was placed on a magnetic rack for 2 minutes or until the supernatant has cleared.
4. The supernatant was removed.
5. On a magnetic rack, the beads were washed with freshly prepared 70% ethanol twice according to manufacturer’s instructions.
6. On a magnetic rack, excess ethanol was removed using aspirator and gel loading tips.
7. On a magnetic rack, the beads were air-dried for <15 minutes and checked in 10 minutes to make sure the beads were not over dry.
8. The sample tubes were removed from the magnetic rack. 30-40 µl of ultra-pure water or elution buffer of choice was added.
9. The solution was gently mixed up and down 10 times to make sure the beads were fully resuspended.
10. The mixture was incubated at room temperature for 10-15 minutes.
11. The tube was placed on the magnetic rack for 2 minutes or until the supernatant has cleared.
12. The supernatant was carefully transferred to a new appropriately labeled 1.5 mL tube.
13. 5 µl of PCR2 products were run on a 2% agarose gel to confirm the band size.
14. The products were frozen at −20°C or −80°C before being submitted to next-generation sequencing facility.

### 13. Next-generation sequencing (NGS)

To select the correct NGS setting, the required read depth can be calculated as below.

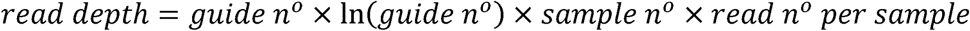

Read depth: the total of reads for all sequenced samples.
Guide n°: the number of guides in the selected library.
Guide n° x ln(guide n°): coupon collector’s expectation, the number of reads required for every guide to be sampled at least once.
Sample n°: the total number of samples across all replicates.
Read n° per sample: the expected reads per sample is typically 300.

Read length is also determined to ensure the regions of interest are covered. For our screen, a read length of 150 cycles was selected with 30% PhiX spike-in. Typically, 10% - 50% PhiX spike-in can be employed to control for sequence clustering and diversity. The percentage of PhiX spike-in to be chosen varies depending on the sequencing platform and should be verified by the sequencing facility. Quality controls including fragment analysis (for example: Fragment analyzer HS NGS assay) and concentration measurement (for example: Qubit HS DNA assay) must be performed, either by the user or the sequencing facility, to confirm library quality prior to sequencing.

The NGS data were next analyzed using the CRISPRCloud2 platform^55^. Briefly, on CRISPRCloud2 site, the Enrichment-based screen option was selected. The kinome library (the reference library) obtained from the manufacturer was converted from excel format to fasta format. Briefly, sequence identifiers and corresponding sequences were organized in Excel, with FASTA headers created by prefixing identifiers with “>” and combining them with sequences using line breaks. The formatted entries were copied from Excel into a plain text editor and saved with a .fasta extension to generate the FASTA file for downstream analysis. High GFP, low GFP, and unsorted/bulk populations of one condition from all replicates were analyzed concurrently. After providing all the necessary information, the web browser initiated the processes of trimming, mapping, and quantifying the sgRNA reads. The processed data were accessed through the link provided. CB^2^ demonstrates several metrics regarding the data quality including gene-level corrected p-values or false-discovery rate, which is a critical factor in selecting hits.

## Discussion

In this paper, we present an optimized kinome-wide screen workflow that facilitates high-throughput screens of several stress-induced autophagy pathways and allows for direct comparison of multiple datasets. While we developed this workflow using an autophagy reporter, the problem it solves, signal drift when analyzing a reporter whose stability is time sensitive, is applicable to other systems where traditional approaches may yield sub-optimal results. Our protocol offers several advantages. Using a fluorescence reporter responsive to a wide variety of stressors allows for the comparison of different pathways in the same cell line. Additionally, the use of a dual fluorescent reporter where one fluorophore acts as a control for reporter expression allows the differentiation pathways regulating transcription and translation versus degradation^17,19^.

Another essential addition to our screen workflow is sample fixation. Our data shows that fixation of cells with PFA prevents the rise in false positives and cell death that comes with long incubation of live cells on ice prior to cell sorting. However, fixation comes with some trade-offs, such as a decrease in read numbers compared to unfixed plates which increased the starting number of cells required for the screen.

Lastly, the CRISPR/Cas9 system used in our study often leads to complete loss of gene function, which can be impractical for studying essential genes or partial loss of functions. However, application of RNAi or CRISPRi approaches can overcome this limitation, especially when emulating pharmacological inhibition^59^.

Collectively, this protocol provides optimizations that aim to greatly reduce the false positive rates of comparative pooled screens. This addresses a general trade-off in multicondition pooled screening where is the fidelity of the sorting is inversely correlated with the number of conditions or library size. These methods can be adapted to other FACS-based pooled screens opening the door to apply high throughput pooled screens to new questions that may have been technically infeasible without this adaptation.

## Abbreviations

BSA: bovine serum albumin
Cas9: CRISPR-associated protein 9
CRISPR: clustered regularly interspaced short palindromic repeats
CRISPRi: CRISPR interference
DMEM: Dulbecco’s modified Eagle medium
DsRed: Discosoma sp. red fluorescent protein
EDTA: ethylenediaminetetraacetic acid
ER: endoplasmic reticulum
FACS: fluorescence-activated cell sorting
gDNA: genomic DNA
GFP: green fluorescent protein
HEK293A: human embryonic kidney 293A cells
HEK293T: human embryonic kidney 293T cells
IRES: internal ribosome entry site
MOI: multiplicity of infection
NGS: next-generation sequencing
p62: sequestosome 1
PBS: phosphate-buffered saline
PCR: polymerase chain reaction
pDNA: plasmid DNA
PEI: polyethylenimine
PFA: paraformaldehyde
PMP70: 70 kDa peroxisomal membrane protein
RETREG1: reticulophagy regulator 1
RNAi: RNA interference
RRID: Research Resource Identifier
SDS: sodium dodecyl sulfate
sgRNA: single guide RNA
TU: transducing units
ULK1: unc-51 like autophagy activating kinase 1

## Acknowledgments

The authors thank the following Core Facilities from the University of Ottawa and the Ottawa Hospital Research Institute (OHRI) for use of their facility, equipment, and expertise: the Flow Cytometry and Virometry Core (RRID:SCR_023306), the Genome Engineering and Molecular Biology Core (RRID:SCR_022954), and the OHRI StemCore Laboratories (RRID:SCR_012601).

## Data availability

All data used in this study are available upon request.

